# A fast and simple method for detecting identity by descent segments in large-scale data

**DOI:** 10.1101/2019.12.12.874685

**Authors:** Ying Zhou, Sharon R. Browning, Brian L. Browning

## Abstract

Segments of identity by descent (IBD) are used in many genetic analyses. We present a method for detecting identical-by-descent haplotype segments that is optimized for large-scale genotype data. Our method, called hap-IBD, combines a compressed representation of genotype data, the positional Burrows-Wheeler transform, and multi-threaded execution to produce very fast analysis times. An attractive feature of hap-IBD is its simplicity: the input parameters clearly and precisely define the IBD segments that are reported, so that program correctness can be confirmed by users.

We evaluate hap-IBD and four state-of-the-art IBD segment detection methods (GERMLINE, iLASH, RaPID, and TRUFFLE) using UK Biobank chromosome 20 data and simulated sequence data. We show that hap-IBD detects IBD segments faster and more accurately than competing methods, and that hap-IBD is the only method that can rapidly and accurately detect short 2-4 cM IBD segments in the full UK Biobank data. Analysis of 485,346 UK Biobank samples using hap-IBD with 12 computational threads detects 231.5 billion autosomal IBD segments with length ≥2 cM in 24.4 hours.

## Introduction

Segments of identity by descent (IBD) are genomic regions over which a pair of individuals share a haplotype due to inheritance from a recent common ancestor. IBD segments are useful in a wide variety of applications because they capture information about genetic relationships between individuals. Correlation between pairwise IBD and phenotypic similarity can be used to detect genomic regions harboring trait-affecting variants^1-6^ and to estimate heritability.^7-10^ IBD segments are also used to estimate kinship coefficients,^11^ detect close relationships,^12-14^ and identify fine-scale population structure.^15-20^

Recent demographic history can be inferred from IBD segments.^15; 16; 21-24^ Populations with smaller effective population size have more IBD sharing because individuals are more closely related on average. Short segments have a larger time to the most recent common ancestor (TMRCA), and thus are informative about less recent effective size, while long segments have a smaller TMRCA and are informative about very recent effective size. Similarly, IBD segments shared between populations are informative about migration rates. Approximately the past 100 generations of demographic history can be inferred from IBD segments.^23^

IBD segments are also useful for estimating population genetic parameters, including mutation rates^25-28^ and recombination rates,^29^ and for detecting regions undergoing recent selection.^10; 20; 30-32^ The mutation rate is estimated from the observed discordance rate in IBD haplotypes. Recombination rate maps can be estimated using the rate of IBD segment endpoints. Selection is detected by looking for genomic regions with higher rates of IBD sharing.

There are several classes of methods for detecting IBD segments. The first class of methods are probabilistic. These methods include PLINK,^2^ Beagle IBD,^33^ and others.^4; 10; 34-39^ For these methods, the unobserved IBD status for a pair (or set) of individuals at a locus takes two (IBD/non-IBD) or more possible states. Typically, a hidden Markov model is used to infer the IBD state at each marker. In the context of a pedigree, with shared haplotypes inherited only through pedigree founders, this IBD-state approach makes sense. However, in population samples, the concept of “IBD state” is ill-defined. Two haplotypes are identical by descent if they are descended from a common ancestor, which is trivially true for all pairs of haplotypes at each position in the genome.

The second class of methods, which includes all the methods presented in this paper, look for long segments of identical-by-state allele sharing either in phased or in unphased genotype data. These identity-by-state (IBS) methods include GERMLINE^40^ and others.^41-44^ In contrast to most of the probabilistic methods, these methods do not dichotomize pairwise haplotypic sharing into “IBD” and “non-IBD”, but instead dichotomize it into “long IBD” and “not long IBD”, which better fits the realities of population-based IBD sharing. Ideally, reported IBD segments should primarily represent IBD from a single common ancestor, rather than a conflation of segments from multiple ancestors, and this is achieved when the length threshold is relatively long.^45^ A drawback of these methods is that the handling of allelic discordances within IBD segments tends to be ad hoc.

For IBS methods, the requirement that two individuals share a haplotype is more stringent than the requirement that the two individuals share at least one allele in their genotypes across a given region. Thus, haplotype-based methods can detect short IBD segments (e.g 2-10 cM in length) with much greater accuracy than genotype-based methods. However, haplotype-based methods can break up a long IBD segment into a sequence of shorter IBD segments if there are haplotype phasing errors in the long IBD segment. For some downstream applications, it is necessary to perform a merging step after detecting IBD segments in order to recover the original long IBD haplotype. On the other hand, genotype-based methods do not require accurately-phased genotype data, and they can detect long segments (>= 15 cM) with high accuracy, which is sufficient for highly-accurate detection of first and second degree relatives.^13^

A third class of methods are those that combine aspects of probabilistic modelling and length-based thresholding on IBS. Typically these methods detect candidate long shared segments, and then form a likelihood ratio for IBD versus non-IBD.^11; 46-48^ These methods tend to be more computationally efficient than the full probabilistic methods, but they cannot analyze large, biobank data sets, unlike some of the purely length-based IBS methods.

Although “identity by descent” implies allelic identity, in fact positions of discordance will be observed. Causes of this discordance include mutation or gene conversion since the common ancestor, and genotype error. Probabilistic methods allow for these discordances via an error term in the modelling, while length-based methods allow for short, infrequent gaps in allele sharing.

The genome-wide average mutation rate in humans (1.3 × 10^−8^ per basepair per meiosis^28^) is similar to the genome-wide average crossover rate (1.2 × 10^−8^ per basepair per meiosis^49^). Consequently, IBD segments from sequence data will each contain an average of approximately one discordance due to mutation.

Gene conversions occur at a rate of 6 × 10^−6^ per basepair per meiosis in humans,^50^ while crossovers occur at a rate of 1.2 × 10^−8^ per basepair per meiosis on average. Thus, an average of around five hundred basepairs per IBD segment will have been subject to gene conversion since the common ancestor. Within a gene conversion tract in an IBD segment, allelic discordance occurs only at positions that were heterozygous in the individual who underwent gene conversion at the locus. Mean heterozygosity in human populations is generally less than or equal to 1 heterozygote per kb.^51^ Thus five hundred basepairs of gene-converted DNA will result in an average of less than 0.5 discordances per IBD segment (depending on the heterozygosity of the population).

Genotype error rates vary greatly across data sets. Data from two recent studies give genotype error rate estimates of 0.008 per Mb per individual in a large SNP array study^52^ and 25 per Mb per individual for single nucleotide variants in a large sequencing study^53^ (error rates estimated as half the discordance between duplicate samples after quality control filtering, multiplied by the average number of called/assayed variants per Mb). Exclusion of rare variants can decrease the genotype error rate,^54^ particularly for sequence data.^53^

With increasingly large data sets, computational issues become significant. The detection of sets of shared haplotypes can be reduced to linear computational complexity, by means of hashing^40; 44^ or by use of the positional Burroughs-Wheeler transform (PBWT).^43^ However, the generation of pairwise IBD segments from these sets scales quadratically with sample size, because the number of pairs of individuals grows quadratically with sample size. Consequently, detecting IBD segments in biobank-scale data is challenging. As well as computation time being an issue, some algorithms require unfeasibly large amounts of computer memory to analyze such data sets.

In this work, we present hap-IBD, which scales to biobank-sized data, provides greater accuracy than competing methods, and is notable for the simplicity of its algorithm and tuning parameters. Hap-IBD utilizes the PBWT^55^ and parallel computation to reduce computing time, and it uses data compression to reduce memory requirements.^56^ It addresses the issue of allele discordance in IBD segments by requiring that a reported segment have a central core (the “seed”) that is free of discordance, while allowing extension beyond the seed after a short gap containing discordance. The key parameters for hap-IBD are the minimum seed length, the minimum extension length, the maximum gap length, and the minimum length of reported IBD segments. These parameters directly control which IBD segments are detected and reported. The hap-IBD program is open-source and freely available for academic and commercial use.

## Methods and Materials

### The hap-IBD algorithm

The hap-IBD method employs a simple seed-and-extend algorithm. A seed is an IBS segment with genetic length greater than a specified minimum value. The hap-IBD algorithm finds all seed segments, and extends each seed if possible. A seed segment is extended if there is another long IBS segment for the same pair of haplotypes that is separated from the seed segment by a short non-IBS gap. The maximum number of base pairs between the first and last markers in the non-IBS gap and the minimum genetic length of the extension IBS segment are specified by the user. A segment may be extended multiple times. When it is no longer possible to extend the segment, the segment is written to the output file if its genetic length is greater than a specified minimum output length.

Allowing short non-IBS gaps provides robustness to three sources of discordant alleles in IBD segments: genotype error, gene conversion, and mutation since the most recent common ancestor. Genotype error and mutation will typically introduce a single discordant allele in an IBD segment. Gene conversion will generally produce a very short interval containing one or a few discordant alleles in an IBD segment. When the phasing of the surrounding alleles is correct, the mis-matching alleles on the pair of IBD haplotypes result in two IBS segments for the same pair of haplotypes, separated by a single marker, or at most a few markers in the case of gene conversion. Our method allows these breaks in IBS sharing to be detected and for the IBS segments on each side of the break to be included in the same reported IBD segment.

Two or more distinct IBS seed segments can result in the same IBD haplotype after each seed is extended. If an IBS segment that extends the seed segment to the left is itself a valid seed segment, we stop the extension process and discard the seed segment that is being extended because the same IBD haplotype will be generated by a seed segment that occurs earlier on the chromosome.

The hap-IBD algorithm also has an optional min-markers parameter that requires seed IBS segments to have a minimum number of markers. The min-markers parameter can be useful for ensuring a minimum level of evidence for IBD in genomic regions having low marker density. When a min-markers parameter is specified, IBS segments that extend a seed are also required to have a minimum number of markers. We set the minimum number of markers in an extension to be the product of the min-markers parameter and the ratio of the minimum extension length to the minimum seed length.

We first describe a single-threaded implementation of the preceding algorithm and then describe how the single-threaded implementation is modified to permit parallel computation.

### Computationally efficient detection of seed segments

After the genotype data for a chromosome are read into memory we apply the positional Burrows-Wheeler transform (PBWT)^55^. The PBWT sweeps through the markers in chromosome order, and at each marker sorts the reverse haplotype prefixes in lexicographic order (the reverse haplotype prefix at the *m*-th marker is the sequence of alleles at markers *m* − 1, *m* − 2, …). At marker *m* we generate a “divergence” array that stores the first marker of the IBS segment containing marker *m* − 1 for each pair of haplotypes that are adjacent after sorting.^55^ The divergence array is used to efficiently identify all seed IBS segments that end at marker *m* (see Durbin’s Algorithm 3).^55^ After a seed is identified, it is extended if possible by comparing the alleles on the two IBS haplotypes in the regions preceding and succeeding the seed segment as described above.

### Memory-efficient computation

The hap-IBD program takes phased genotype data in VCF format as input.^57^ As the genotype data are read into memory, the data are immediately converted to binary reference format (version 3).^56^ Binary reference format compresses low frequency variants by storing only the indices of the haplotypes carrying non-major alleles. Higher frequency variants are compressed by storing unique allele sequences in a region, along with a vector that maps haplotypes indices to the allele sequence carried by the haplotype. We use binary reference format because it permits data for an entire chromosome to be stored compactly in memory, and it allows rapid queries of alleles carried by haplotypes at each marker.

The PBWT requires only two additional arrays of stored information, each with length equal to the number of haplotypes. Seed IBS segments are extended as soon as they are identified by the PBWT. After extension, segments that are longer than the minimum output segment length are immediately printed to an output buffer, which is flushed to disk when full. Consequently, only a limited number of IBD segments are stored in memory at any time.

### Parallelization

The hap-IBD algorithm is parallelized by applying the PBWT concurrently in overlapping marker windows. If *L* is the genetic distance between the first and last markers on the chromosome, *S* is the minimum seed genetic length, and *T* is the number of computational threads, we sequentially define *T* overlapping marker windows *W*_1,_ *W*_2,_ … *W*_*T*_ that each have length approximately equal to ((*L* − *S*)/*T* + *S*) cM, and that have approximately *S* cM overlap between adjacent windows. The first window *W*_1_ begins at the first marker on the chromosome and ends at the first marker after genetic position ((*L* − *S*)/*T* + *S*) whose index is greater than the minimum number of markers required for an IBS seed segment. The first marker in *W*_*i*+1_ is the first marker in *W*_*i*_ that cannot be the start of a seed IBS segment contained within *W*_*i*_ because the number of markers or genetic distance separating the marker from the last marker in *W*_*i*_ is too small. The last marker in *W*_*i*+1_ is the first marker that is ≥ (*L* − *S*)/*T* cM away from the last marker in window *W*_*i*_. With these definitions every seed IBS segment will be detected in at least one of the overlapping windows.

We run the PBWT algorithm in each overlapping window in parallel. When a seed IBS segment is found, we ignore the window boundaries when we extend the segment, so that the extension process is the same as for the single-threaded case. If multiple seeds result in the same maximal IBD segment after extension, we keep the maximal IBD segment generated by the first seed, and discard the duplicate IBD segments generated by later windows seeds.

### Input and output data

The input data is a VCF file^57^ with phased, non-missing genotype data, and a PLINK-format genetic map.^2^ Linear interpolation is used to estimate the genetic map positions for any marker whose position is not on the genetic map. Although the use of a genetic map is recommended, hap-IBD can be used with Mb units simply by supplying a genetic map with a recombination rate of 1 cM = 1 Mb.

Two output files are produced: one containing within-individual segments of homozygosity by descent (HBD) and one containing between-individual IBD segments. Each output line contains the pair of samples, the specific haplotypes, the starting base position, the ending base position, and the genetic length of the HBD or IBD segment.

### Analysis parameters

The minimum seed length parameter has a large influence on computation time. Increasing the minimum seed length reduces computation time because fewer seed IBS segments will be considered. Decreasing the minimum seed length can increase power to detect short IBD segments that have discordant alleles on the pair of shared haplotypes.

The maximum gap length and minimum extension length allow reported IBD segments to contain discordant alleles due to genotype error, mutation, or gene conversion. The hap-IBD software also has an option for excluding input markers having low minor allele count.

The minimum markers parameter controls the minimum number of markers in IBS seed and extension segments. The number of reported IBD segments should be approximately constant throughout the genome; however, regions with low marker density can produce local spikes in the number of reported IBD segments (see Results). These spikes contain many IBS segments that satisfy the genetic length requirements, but which contain relatively few markers. The spikes can be reduced or eliminated by post-processing,^23; 58^ or by requiring seed and extension IBS segments to contain a minimum number of markers.

### UK Biobank genotype data

We downloaded the UK Biobank genotype data from the European Genome-phenome Archive^59^ (Dataset accession: EGAD00010001497). The UK Biobank data contain 488,377 individuals and 784,256 autosomal markers.^52^ We excluded markers with more than 5% missing genotypes (n = 70,246), markers that had only one individual carrying a minor allele (n = 5,123), and markers that failed one or more of the UK Biobank’s batch quality control tests (n = 1,527).^52^ After excluding 72,601 markers that failed one or more of these filters, there were 711,655 autosomal markers.

We then exclude 968 individuals that were identified by the UK Biobank as being outliers for their proportion of missing genotypes or proportion of heterozygous genotypes, and we excluded 9 individuals that were identified by the UK Biobank as showing third degree or closer relationships with more than 200 individuals (indicating sample contamination).^52^ After these exclusions there were 487,400 individuals.

We identified parent-offspring trios using the kinship coefficients and the proportion of markers that share no alleles (IBS0) that are reported by the UK Biobank.^52; 60^ First degree relatives were considered to be pairs of individuals with kinship coefficient between 2^−2.5^ and 2^−1.5^. Among first degree relatives, parent-offspring relationships were assumed to be the first-degree relative pairs with IBS0 < 0.0012. These are the same kinship coefficient and IBS0 thresholds used by the UK Biobank to identify parent-offspring relationships.^52^ We considered an individual to be the offspring in a parent-offspring trio if the individual had a parent-offspring relationship with exactly one male and one female individual, and if the male and female first-degree relatives are not in the set of related pairs of individuals reported by the UK Biobank, which is the set of pairs of individuals with estimated kinship coefficient greater than 2^−4.5^. In this case, we considered the male and female first-degree relatives to be the offspring’s parents. Using this procedure, we identified 1064 parent-offspring trios.

The 1064 trio offspring have 2,054 distinct parents. We excluded these parents from the data before phasing and IBD segment detection so that phasing accuracy in the trio offspring would more closely match phasing accuracy in unrelated individuals. After excluding the trio parents, there were 485,346 remaining individuals. We listed the 1064 trio offspring followed by the remaining samples in random order. We created five telescoping genotype data sets that included 5000, 15,000, 50,000, 150,000, and all 485,346 individuals by taking the corresponding number of samples from the top of this list. We then phased each data set with Beagle 5.1.^61^

We used the parental genotype data to determine true phase in 850 trio offspring. We selected the 850 trio offspring by computing the number of autosomal sites with Mendelian inconsistent genotypes in each of the 1064 trios (range: 57-5102 inconsistencies) and taking the offspring from the trios having the smallest number of Mendelian inconsistent genotypes (range: 57-456 inconsistencies).^62^ We phased the 850 trio offspring at all heterozygous genotypes for which phase could be determined from parental genotypes and Mendelian inheritance constraints (82.4% of heterozygous genotypes), and we masked genotypes at Mendelian inconsistent sites in this phased data. We used these estimated haplotypes to evaluate false-positive and false-negative rates for IBD segment detection as described below.

After excluding trio parents, there were 43 remaining parent-offspring pairs who were not part of a trio in the 50,000 individual subset of the UK Biobank data. We use these 43 remaining pairs to compute the mean proportion of chromosome 20 covered by detected IBD segments in parent-offspring pairs.

### Simulated Data

In order to test the performance of hap-IBD and other methods on sequence data, we generated 60 Mb of data for 50,000 individuals from a demographic model that simulates the present UK European population.^47^ This model has a population size of 24,000 in the distant past, a reduction to 3,000 occurring 5,000 generations ago, growth at rate 1.4% per generation starting 300 generations ago, and growth at rate 25% beginning ten generations ago.

We used forward simulation with SLiM^63^ to simulate the ancestral recombination graph for the most recent 5000 generations. Gene conversion tracts were initiated at a rate of 2 × 10^−8^ per bp per generation, and had geometrically distributed lengths with mean 300 bp, giving an overall gene conversion rate of 6 × 10^−6^. ^27; 50^ A constant recombination rate of 1 × 10^−8^ was used. We then used msprime’s coalescent simulation to add mutations (at rate 1.38 × 10^−8^) and simulate the more distant past.^64^ This hybrid strategy of using SLiM and msprime enables utilization of msprime’s computational efficiency for large data sets, while incorporating biologically realistic settings such as gene conversion that are implemented in SLiM but not currently implemented in msprime.^65^ Our simulation only includes gene conversion events in the most recent 5000 generations, but it is the more recent gene conversions that have the greatest potential impact on haplotype phase accuracy and that can create discordances between identical by descent haplotypes.

We determined the true IBD segments for 1000 simulated individuals from the simulated ancestral recombination graphs. IBD segments are required to have the same ancestral node along their length, except for short breaks due to gene conversion.

We added genotype error at a rate of 0.02%, which is the error rate that produces the observed 0.04% rate of discordance at SNVs passing quality control in the TOPMed Freeze 5 whole genome sequence data.^53^ We then removed variants with frequency less than 0.10 and phased the remaining genotypes using Beagle v5.1.^61^ We also separately phased a subset of 5000 individuals with the same minor allele frequency threshold of 0.10. We found that phasing accuracy for common variants improves when rare variants are excluded, and that the improved phasing accuracy more than offsets the loss of information from excluding low frequency variants. Low frequency variants are not very informative for IBD because most individuals are homozygous for the major allele, and because allele discordance at low frequency variants in IBD segments could due to genotype error, recent mutation, or phasing error, rather than indicating non-IBD. Other methods for IBD detection in sequence data have used a minor allele frequency filter. The application of GERMLINE to the Genomes of the Netherlands whole genome sequence data used a minor allele frequency filter of 1%.^58^ The TRUFFLE analysis of 1000 Genomes sequence data used minor allele frequency filters of 5% and 10%.^42^

### Parameter settings

Each method has an option for setting the minimum length of reported IBD segments. All methods, except TRUFFLE, measure genetic distance in cM units. For TRUFFLE, we substituted Mb units for cM units. All genetic distances are interpolated from the HapMap genetic map.^66^

We required all UK Biobank chromosome 20 analyses to complete within two days of wall-clock time on the compute nodes used for these analyses. We set the minimum output segment length to 2.0 cM, unless a higher output segment length was required for analyses to finish within two days of wall-clock time. Parameter settings for analysis of UK Biobank and simulated sequence data are based on previously published analyses of SNP array^5; 42-44^ and sequence data.^42; 43; 58^ Parameter settings for each method are reported in Tables S1 and S2. We do not evaluate iLASH on the sequence data because the published description of this method does not include analyses of sequence data.

### Comparison of methods

For the simulated data, coalescent trees for 1000 simulated samples were used to determine true IBD segments exceeding 1.5 cM in length for those samples. For the UK Biobank data, we considered true IBD segments to be IBS segments exceeding 1.5 cM in length among the 850 trio offspring which were phased using parental genotypes and Mendelian inheritance rules.

#### False-positive rate estimation

We divided detected IBD segments into bins according to the detected segment length (2-3, 3-4, 4-6, 6-10, 10-18, and >18 cM). For each detected IBD segment, we identified the cM length of the portion of the detected segment that is not covered by any true IBD segment with length >1.5 cM, and we calculated the sum of these false-positive segment lengths. The false-positive rate for a bin is the sum of the false-positive segment lengths divided by the sum of the detected segments lengths.

#### False-negative rate estimation

We divided true IBD segments into bins according to the true segment length. The length bins and number of true IBD segments in each length bin are: 2.5-3 cM (2492), 3-4 cM (1360), 4-6 cM (551), 6-10 cM (160), 10-18 cM (55), and >18 cM (64). For each true IBD segment, we identified the cM length of the portion of the true segment that is not covered by any detected IBD segment with length >2.0 cM, and we calculated the sum of these false-negative segment lengths. The false-negative rate for the bin is the sum of the false-negative segment lengths divided by the sum of the true segments lengths.

#### ROC analysis

In order to account for inter-method differences in determining IBD end-points, which affect the reported length of IBD segments, we calculated false positive and false negative rates for each method over a range of detected segment length thresholds (1.6, 1.7, 1.8, 1.9, 2.0, 2.1, 2.2, 2.3, and 2.4 cM). We calculated the false positive rate for each threshold as described above using all true segments having length ≥ 1.5 cM, and we calculated the false negative rate as described above using all true segments having length ≥ 2.5 cM. We then generated a receiver operating characteristic (ROC) curve for each method that shows the true-positive rate (which is one minus the false-negative rate) and false-positive rate for each detected segment length threshold. For length thresholds < 2.0 cM, the hap-IBD minimum seed length was set to 1.6.

We also generated ROC curves for each method for 5 cM segments. For this analysis, we used detected segment length thresholds of 4.6, 4.8, 5.0, 5.2, and 5.4. We calculated false positive rates using all true segments having length ≥ 4.5 cM, and we calculated false negative rates using all true segments having length ≥ 5.5 cM.

#### Computation time

All analyses were run on a 12-core 2.6 GHz computer with Intel Xeon ES-2630 processors and 128 GB of memory. Computation time was measured using the unix time command, which returns a “real”, a “system”, and a “user” time. The wall-clock time is the “real” time, which is the length of time the program was running. The CPU time is the sum of the “system” and “user” times. For multi-threaded compute jobs, the CPU time includes the sum of the CPU time for each computational thread, so that it represents the total CPU resources consumed by the program. A maximum of 2 days of wall-clock time was allowed for each analysis of UK Biobank data chromosome 20 data or of the 60 Mb of simulated sequence data, with no results reported if the analysis did not complete within this time frame.

## Results

### Computational feasibility

Figure 1 shows CPU times for subsets of the UK Biobank chromosome 20 data (5000 to 485,346 individuals) for 2 cM and 5 cM output length thresholds. For the full data UK Biobank data with 485,346 individuals, hap-IBD detected 3.43 billion IBD segments on chromosome 20 at the 2 cM threshold, and 106 million segments at the 5 cM threshold. GERMLINE could not analyze subsets of size 50,000 or more individuals because it required more than 128 Gb of memory. TRUFFLE could not analyze subsets of size 150,000 or more individuals on our compute nodes. iLASH could not analyze subsets of 150,000 or more individuals at the 2 cM output threshold because it needed more than 128 Gb of memory, but it could analyze the full data set at the 5 cM output threshold. RaPID could not analyze the full chromosome 20 data set at the 2 cM threshold within the permitted two days of wall-clock time, but it could analyze the full data set at the 5 cM output threshold.

**Figure 1.**
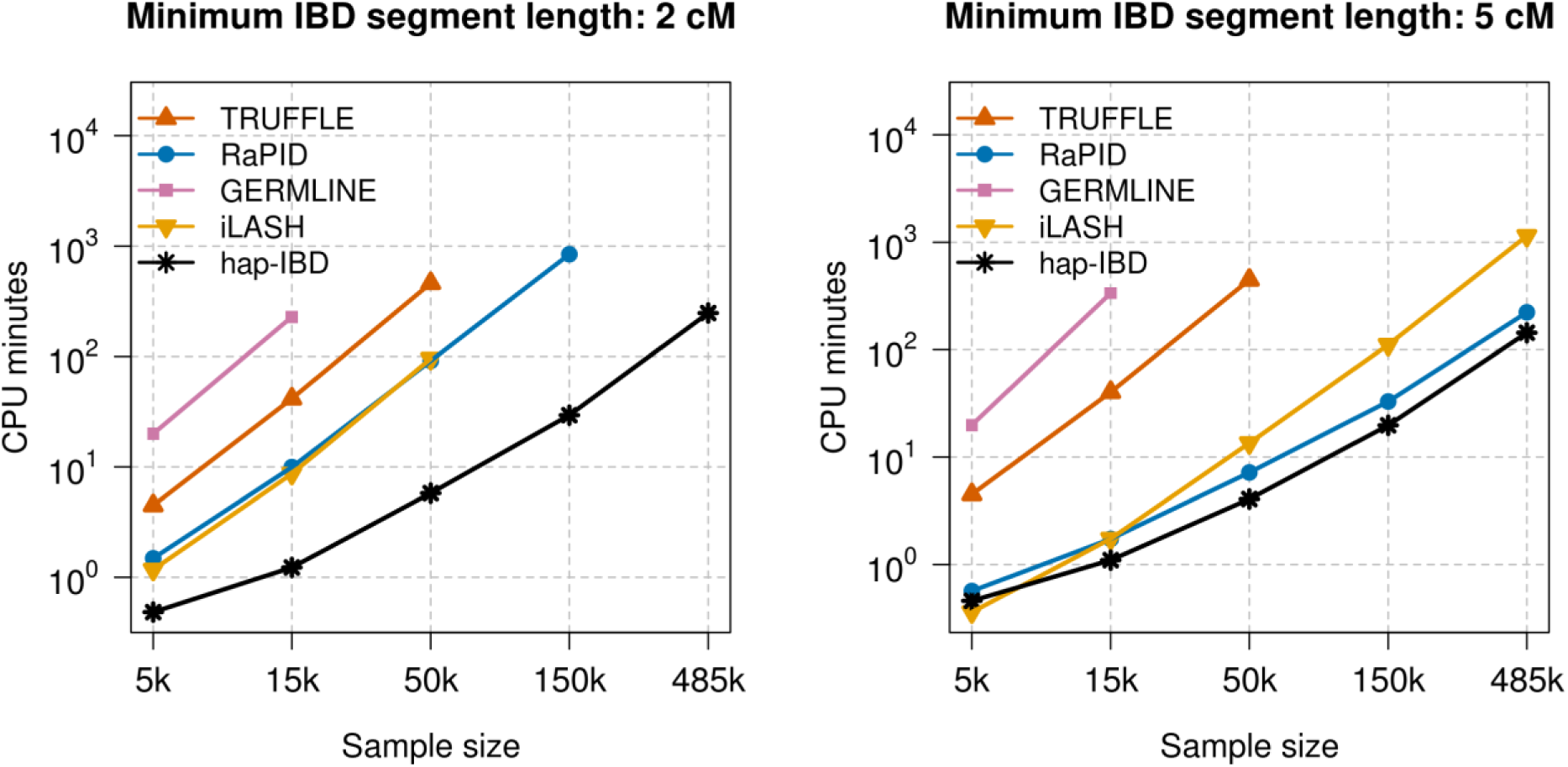
CPU time. CPU time for detecting IBD segments with length ≥ 2 cM (left panel) and ≥ 5 cM (right panel) on chromosome 20 in samples of 5000, 15,000, 50,000, 150,000, and 485,346 individuals from the UK Biobank. CPU time is the sum of the computation time for each CPU core. All programs used 12 computational threads, except RaPID and GERMLINE which are limited to one computational thread.

Three of the five methods have parallelization implemented, and running these methods on a 12-core computer leads to an approximate 10-fold reduction in computing time compared to single-threaded analysis (Figure S1). This degree of speedup is important for analysis of large data sets. For example, the single-threaded RaPID program required 223.6 minutes of wall-clock time to output IBD segments > 5 cM for all samples on chromosome 20, but hap-IBD required only 13.4 minutes when using 12 computational threads.

Overall, we see that hap-IBD is the fastest program except when analyzing the smallest sample size (5000 individuals) using the largest output threshold (5 cM output threshold); for this combination iLASH is faster. In our experiments, hap-IBD was the only method that could analyze the full UK biobank chromosome 20 data on our compute servers in less than 2 days when using a 2 cM output length threshold.

We also performed a genome-wide analysis of the 22 autosomes for the UK Biobank data. Analysis of 485,346 UK Biobank samples using hap-IBD with 12 computational threads detected 231.5 billion autosomal IBD segments in 24.4 hours.

### Accuracy

Several methods have a low false-positive rate (Figures 2 and S2) but a high false-negative rate (Figures 3 and S3), or vice versa. The methods apply different algorithms for determining the end-points of IBD segments, and this results in different methods reporting different lengths for a true IBD segment. Since false-positive rates and false-negative rates can be traded off by changing the output length threshold, we constructed ROC curves by varying the output IBD segment length threshold for each method in order to assess the true-positive vs false-positive trade-off. The true-positive rate is one minus the false-negative rate. An ideal method would have a true-positive rate of 1 and a false-positive rate of 0. For 2 cM IBD segments (Figure 4) and for 5 cM IBD segments (Figure S4) hap-IBD shows the best performance on these ROC curves. In particular, hap-IBD has much lower false-positive rates than RaPID and much higher true-positive rates than iLASH. The IBD segment detection method for unphased genotype data (TRUFFLE) has high error rates for these short IBD segments.

**Figure 2.**
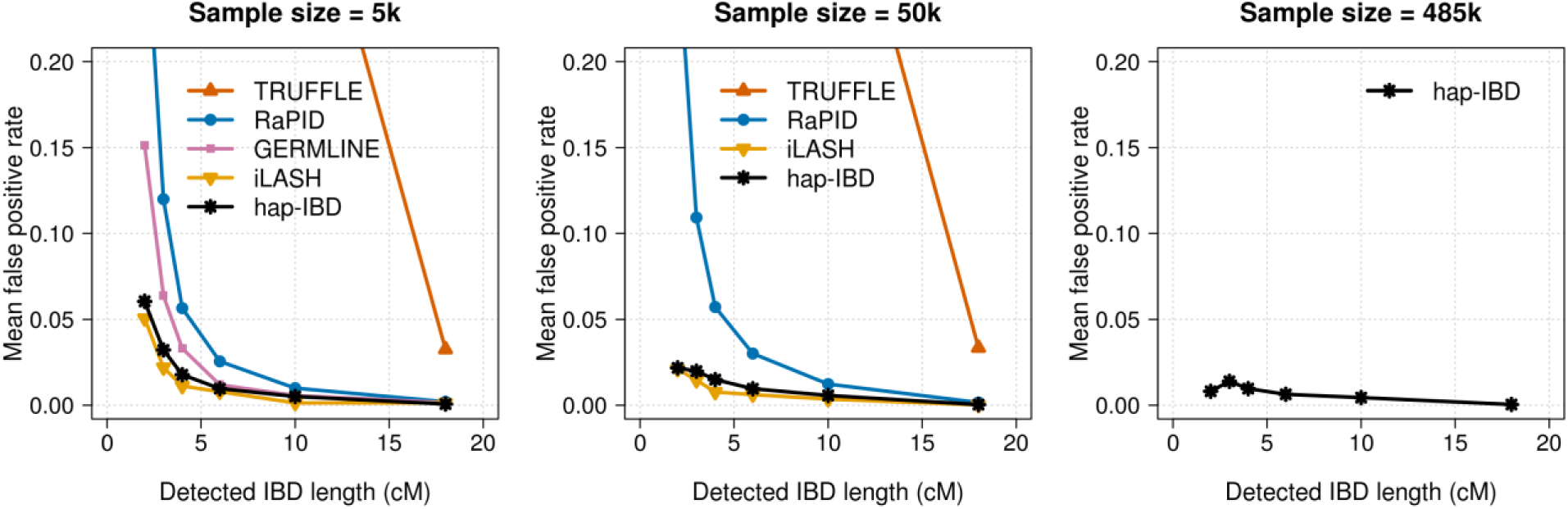
False-positive IBD segment detection in UK Biobank chromosome 20 data. False-positive rates for IBD segment detection for 5000, 50,000, and 485,346 UK Biobank samples. IBD segments with length ≥ 2 cM were detected with each method. Detected IBD segments were assigned into bins of 2-3, 3-4, 4-6, 6-10, 10-18, and >18 cM according to their segment length. The false-positive rate is the proportion of detected IBD segments in a bin that are not covered by any true IBD segment ≥ 1.5 cM in length. Hap-IBD is the only method shown for the full UK Biobank analysis (485,346 individuals) because other methods were unable to complete the analysis with a 2 cM output threshold within the memory and time constraints (see Computational Feasibility Results). The x-coordinate of each data point is the left bin end point (e.g. 2 cM for the 2-3 cM bin). For the full range of y-coordinate values see Figure S2.

**Figure 3.**
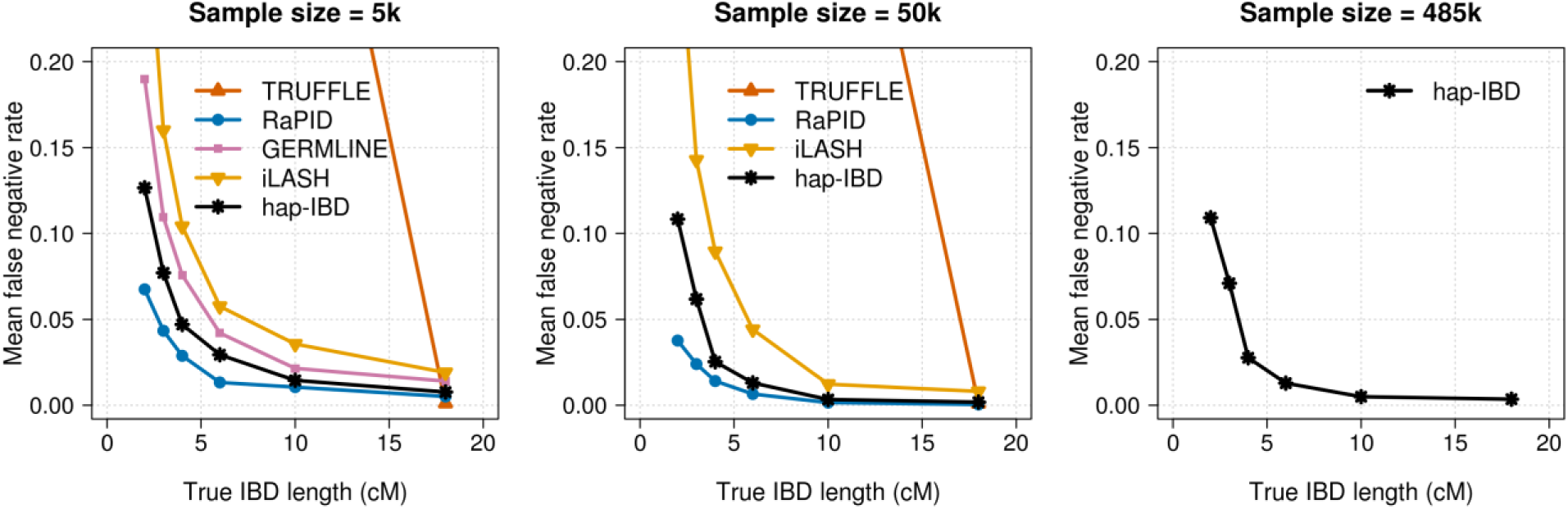
False-negative IBD segment detection in UK Biobank chromosome 20 data. False-negative rates for IBD segment detection for 5000, 50,000, and 485,346 UK Biobank samples. IBD segments with length ≥ 2 cM were detected with each method. The right column of plots shows false negative rates. True IBD segments with length ≥ 2.5 cM were assigned into bins of 2.5-3, 3-4, 4-6, 6-10, 10-18, and >18 cM according to their segment length. The false-negative rate is the proportion of true IBD segments in a bin that are not covered by any detected IBD segment ≥ 2 cM in length. Hap-IBD is the only method shown for the full UK Biobank analysis (485,346 individuals) because other methods were unable to complete the analysis with a 2 cM output threshold within the memory and time constraints (see Computational Feasibility Results). The x-coordinate of each data point is the left bin end point (e.g. 2.5 cM for the 2.5-3 cM bin). For the full range of y-coordinate values see Figure S3.

**Figure 4.**
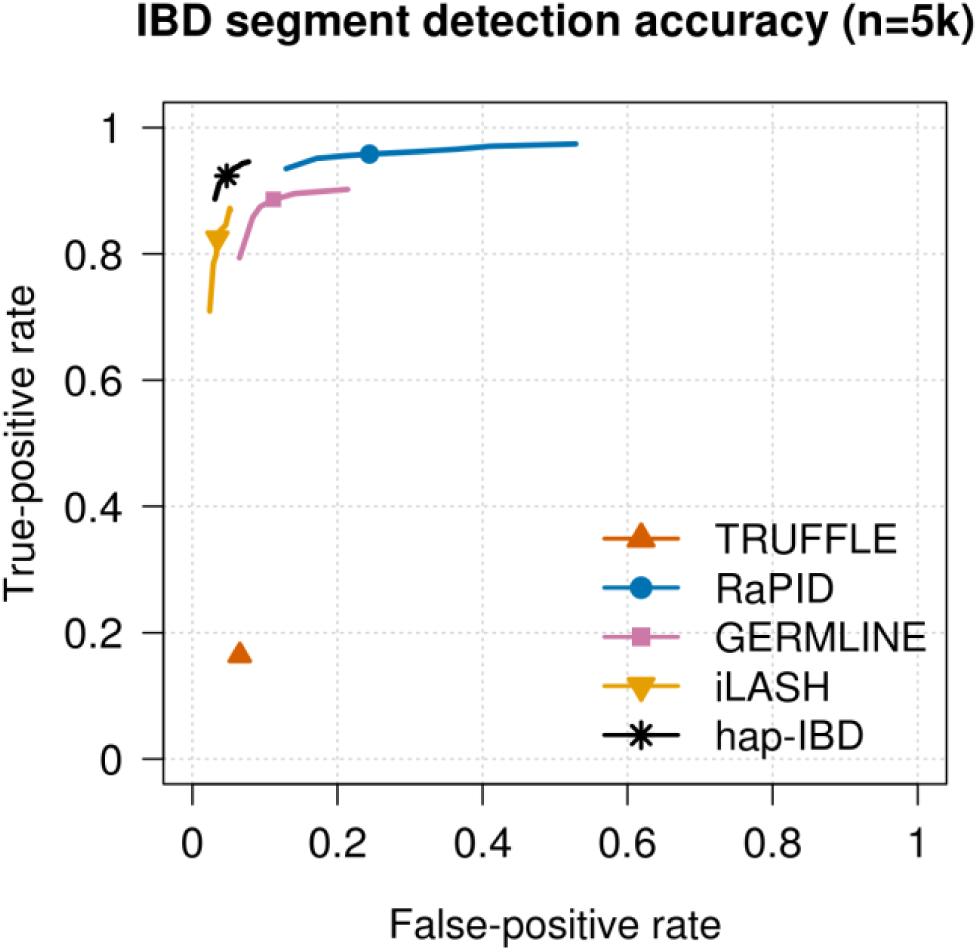
ROC curves for 2 cM IBD segment detection in UK Biobank chromosome 20 data. False-positive and false-negative rates for detection of IBD segments over a range of output length thresholds around 2 cM for 5000 UK Biobank samples. False positives are assessed using true segments of length ≥ 1.5 cM, and false negatives are assessed using true segments of length ≥ 2.5 cM in order to allow for some discrepancy between reported and true lengths. IBD segments were detected with each method using length thresholds of 2 cM (plotted symbol) and with other thresholds between 1.6 and 2.4 cM (lines; see Methods). Figure S4 shows a similar plot for 5 cM.

We also investigated the proportion of chromosome 20 in parent-offspring pairs that was covered by detected IBD segments with length ≥ 2 cM in 43 parent-offspring pairs in the set of 50,000 UK Biobank samples. The proportions were 0.978 for iLASH, 0.987 for hap-IBD, 0.994 for RaPID, and 1.0 for TRUFFLE. GERMLINE was not evaluated because it could not analyze 50,000 individuals on our compute server. All methods detected IBD across all or nearly all of the chromosome in the parent-offspring pairs. For haplotype-based methods, the methods with higher false-positive rates (Figure 2) detected slightly higher amounts of IBD in the parent-offspring pairs. Genotype-based methods are not affected by haplotype phase errors and the genotype-based method (TRUFFLE) had the highest detection rate for these chromosome-length shared haplotypes.

In genome-wide analysis of the UK Biobank data, we find regions in which IBD detection methods report inflated levels of IBD segments. These are generally regions with large gaps in marker coverage, or very low marker density, and often occur around centromeres. Figure 5 shows results for chromosomes 1 and 20 for the methods with the highest accuracy for short IBD segments (the four haplotype-based methods), for the 5000 individual UK Biobank data. Around the chromosome 1 centromere the methods are finding IBD segments at a rate 40 to 3000 times greater than the baseline level. The inflation is worse for RaPID and iLASH than for GERMLINE and hap-IBD. Figure S5 shows that the inflated detection can be reduced by increasing hap-IBD’s min-markers parameter. However, the use of overly high values of this parameter will reduce power to detect short IBD segments. Alternatively, regions with high rates of IBD segment discovery can be identified after IBD segment detection and excluded.^67^

**Figure 5:**
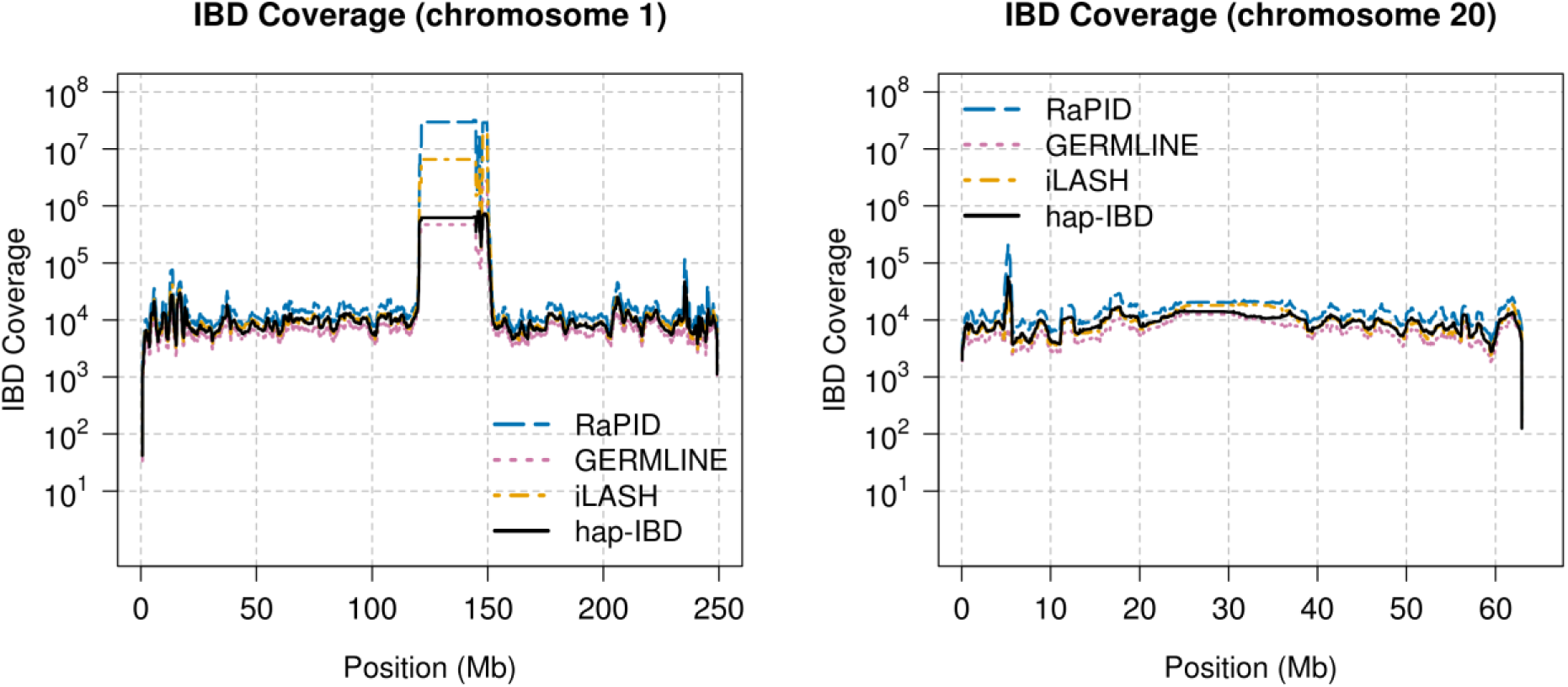
Chromosome-wide IBD segment detection in UK Biobank. The methods were run on 5000 UK Biobank samples on chromosome 1 and chromosome 20 to detect IBD segments of length ≥ 2 cM. Each chromosome is divided into non-overlapping 10 kb intervals. For each interval, the IBD segments intersecting the interval are weighted by the proportion of the 10 kb interval that is covered by the IBD segment, and the sum of weights is plotted as the IBD coverage.

We also assessed accuracy using simulated sequence data. There are several important differences between the UK Biobank analysis and the simulated sequence data analysis. First, the approach to assessing accuracy differs. In the UK Biobank, we determine true phase of trio offspring and use that to determine identity by state at the haplotype level, which we use as a proxy for true IBD. The genotype error rate is extremely low in these data (with a duplicate discordance rate of 6.7 × 10^−5^),^52^ but genotype errors can disrupt both the true IBD and the estimated IBD in the UK Biobank analysis. In contrast, in the simulated data the true IBD status is obtained directly from the simulation (defined as no change in common ancestor across a segment except in tracts of gene conversion), and mis-called alleles may disrupt the estimated IBD but do not affect the ascertainment of true IBD. Second, the marker density is much higher for the simulated sequence data. Although we remove markers with minor allele frequency < 0.1 (see Methods), the marker density is still five times greater than that of the UK Biobank (97,890 markers with minor allele frequency ≥ 0.1 in the simulated 60 Mb region, compared with 18,424 total UK Biobank markers on chromosome 20). Third, the genotype error rate in the simulated sequence data is much higher than for the UK Biobank data. With current technology, error rates tend to be higher for sequence data than for SNP array data, even with high sequence coverage and careful processing. We added genotype error to the simulated sequence data at a rate that generates the level of duplicate discordance observed in the TOPMed data, which is 4 × 10^−4^ for SNPs passing quality control.^53^ This level of duplicate discordance is six times higher than for the UK Biobank SNP data. There are also important similarities between the two analyses, which include the length of the region (approximately 60 Mb for the simulated analysis and for the UK Biobank chromosome 20 analysis), large sample size (up to 50,000 for the simulated data and up to 485,346 for the UK Biobank data), and demographic history (UK-like simulation versus actual UK population).

In the simulated sequence data, we compared methods for which the authors have published analyses of sequence data (GERMLINE, RaPID, and TRUFFLE), and we replicated settings from those published analyses (see Methods for details). To compare accuracy, we produced ROC curves for detection of 2 cM segments. We considered sample sizes of 5000 (Figure 6) and 50,000 (Figure S6). We find that hap-IBD and GERMLINE have a very similar accuracy profile (for the 5000 individuals only, because GERMLINE could not analyze the 50,000 individuals with the available computer memory). TRUFFLE had very low power to detect 2 cM IBD segments, while RaPID had a high false positive rate. Overall these results are similar to those seen in the UK Biobank analysis, except that the relative accuracy of GERMLINE is improved in these simulated sequence data. The parameters that we using for GERMLINE in the simulated sequence analysis may be a better match for these data than were the parameters that we used for the UK Biobank data, although we used published parameter settings in both instances.

**Figure 6.**
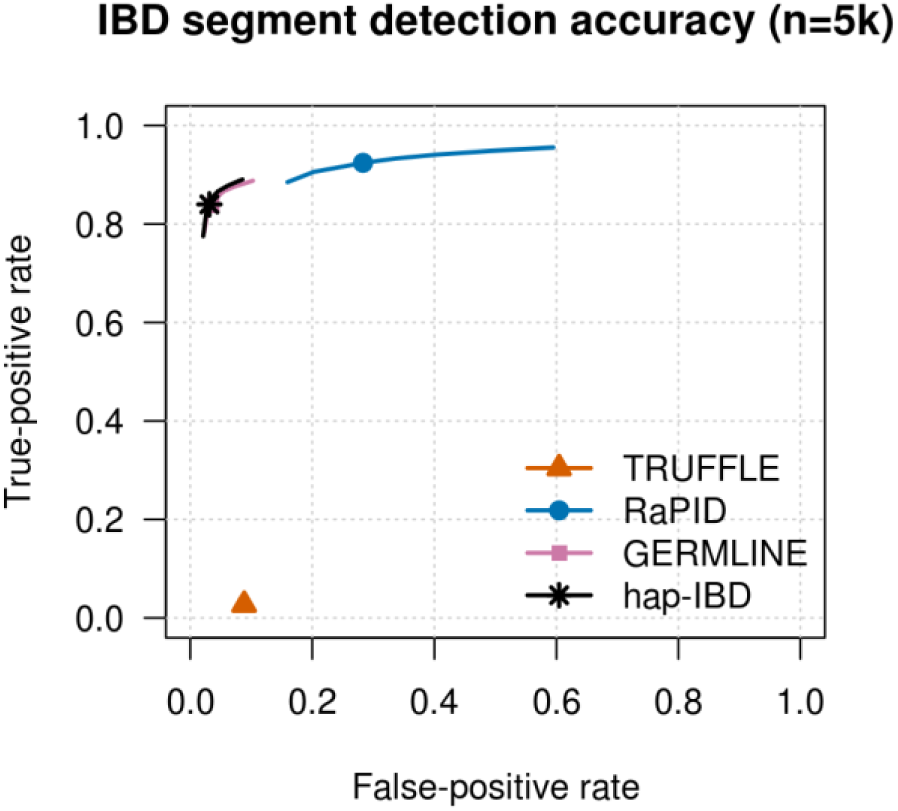
ROC curves for IBD segment detection in simulated sequence data. False-positive and false-negative rates for detection of IBD segments over a range of output length thresholds around 2 cM for 5000 simulated samples. False positives are assessed using true segments of length ≥ 1.5 cM, and false negatives are assessed using true segments of length ≥ 2.5 cM in order to allow for some discrepancy between reported and true lengths. IBD segments were detected with each method using length thresholds of 2 cM (plotted symbol) and with other thresholds between 1.6 and 2.4 cM (lines; see Methods). Figure S6 shows a similar plot for 50,000 samples.

## Discussion

We have presented an IBD segment detection method for large-scale genotype data that is substantially faster and more accurate than four state-of-the-art competing methods (GERMLINE, iLASH, RaPID, and TRUFFLE). We applied hap-IBD to 485,346 samples from the UK Biobank^52^ and detected 231.5 billion autosomal IBD segments having length >2 cM in less than 24.4 hours of wall-clock time on compute server with 12 CPU cores.

An attractive feature of hap-IBD is its simplicity. All seed IBS segments that exceed a specified length are identified and then extended if possible. The extension process allows for sporadic non-IBS alleles due to mutation, genotype error, or gene conversion. The hap-IBD parameters define the minimum length of IBS seed and extension segments and the maximum length of non-IBS gaps. These parameters have a simple and direct relationship to the IBD segments that are reported, which enables the correctness of the results to be confirmed. In contrast, some methods utilize a large number of tuning parameters which have only an indirect relationship to output IBD segments, such as iLASH’s seven parameters for controlling locality-sensitive hashing: perm_count, shingle_size, shingle_overlap, bucket_count, match_threshold, interest_threshold, and minhash_threshold.^44^

The hap-IBD method shares some similarities with the GERMLINE method: both methods search for long IBS segments via a seed and extend algorithm and both methods allow for the presence of some discordant alleles in a reported IBD segment.^40^ However, hap-IBD achieves much greater computational efficiency and greater accuracy than GERMLINE by employing the positional Burrows-Wheeler transform instead of a hash table and by identifying seeds that exceed a specific genetic length rather than a specified number of markers.

In our tests, hap-IBD consistently required less CPU time than competing methods. The hap-IBD method includes internal parallelization that can yield wall-clock compute times that are a fraction of the total CPU time on multi-core processors.

The hap-IBD method requires phased genotype data. In practice, nearly all large genotype data sets are phased because phased data are required to obtain the highest accuracy for many downstream analyses including IBD segment detection, relationship inference,^13^ local ancestry inference,^68^ population demography inference,^21-23^ and detection of selection.^10^ Phased data are also required for computationally efficient and accurate genotype imputation.^56; 69^ With state-of-the-art methods, the effort and computational cost required to phase large data sets is modest when using a small compute cluster. We phased the UK Biobank genome-wide data with Beagle 5.1 in less than two days using 16 compute servers, each with 20 CPU cores.

Our results confirm that IBD segment detection methods for phased genetic data can detect much shorter IBD segments than methods for unphased genetic data. In our tests, the method for unphased data could not accurately detect segments with length < 10 cM, but most methods for phased data could accurately detect IBD segments with length > 2 cM (Figure 4). Furthermore, haplotype-based methods identify the shared allele sequence, whereas genotype methods cannot identify the shared allele when the individuals both carry an identical heterozygote genotype. However, genotype-based IBD detection methods have some advantages. Genotype-based methods can detect first and second degree relationships with high accuracy before haplotypes are estimated,^13; 42^ which can be useful during initial data quality control. In addition, genotype-based methods can be used when genotype data cannot be accurately phased due to non-uniform marker coverage or high rates of genotype error, such as can be the case for exome and low-coverage sequence data.

The hap-IBD method performs well across a range of haplotype switch error rates. In the UK Biobank data, the switch error rate for 5000 samples is more than an order of magnitude higher than the switch error rate for 485,346 samples.^62^ However, even for the 5000 sample subset of the UK Biobank, the IBD-detection accuracy is very high and is sufficient to identify close relatives in the data. Furthermore, one can increase the accuracy of phase estimates in small samples by phasing the samples together with a reference panel of sequenced individuals.^70^

A general limitation of IBD detection segment methods that rely on IBS is that there is some degree of error in determination of segment end points. The IBS interval can extend beyond the end-points of a contained IBD segment. Consequently, IBD detection methods that report the full IBS interval will often over-extend the IBD segment ends. Such methods can also miss some regions at the end of IBD segments when genotype error, mutation, or gene conversion near the end of the IBD segment causes the IBS segment to end before the actual end of the IBD segment. If the genetic distance between the truncated end of the IBS segment and the true end of the IBD region is short, it is not possible determine with confidence whether or not the IBD segment extends past the end of the IBS segment. Development of IBD segment detection methods that are robust to genotype error, recent mutation, and gene conversion that occur near the ends of IBD segments is an area for future research.

## Supplemental Data

Supplemental Data include 2 tables and 6 figures.

## Acknowledgements

Research reported in this publication was supported by the National Human Genome Research Institute and National Institute of General Medical Sciences of the National Institutes of Health under award numbers HG008359, HG005701, and GM075091. The content is solely the responsibility of the authors and does not necessarily represent the official views of the National Institutes of Health. This research has been conducted using the UK Biobank Resource under Application Number 19934.

## Web resources

hap-IBD:

https://github.com/browning-lab/hap-ibd

Beagle,

http://faculty.washington.edu/browning/beagle/beagle.html

The UK Biobank data,

https://www.ebi.ac.uk/ega/home

## Supplementary Information

**Figure S1.**
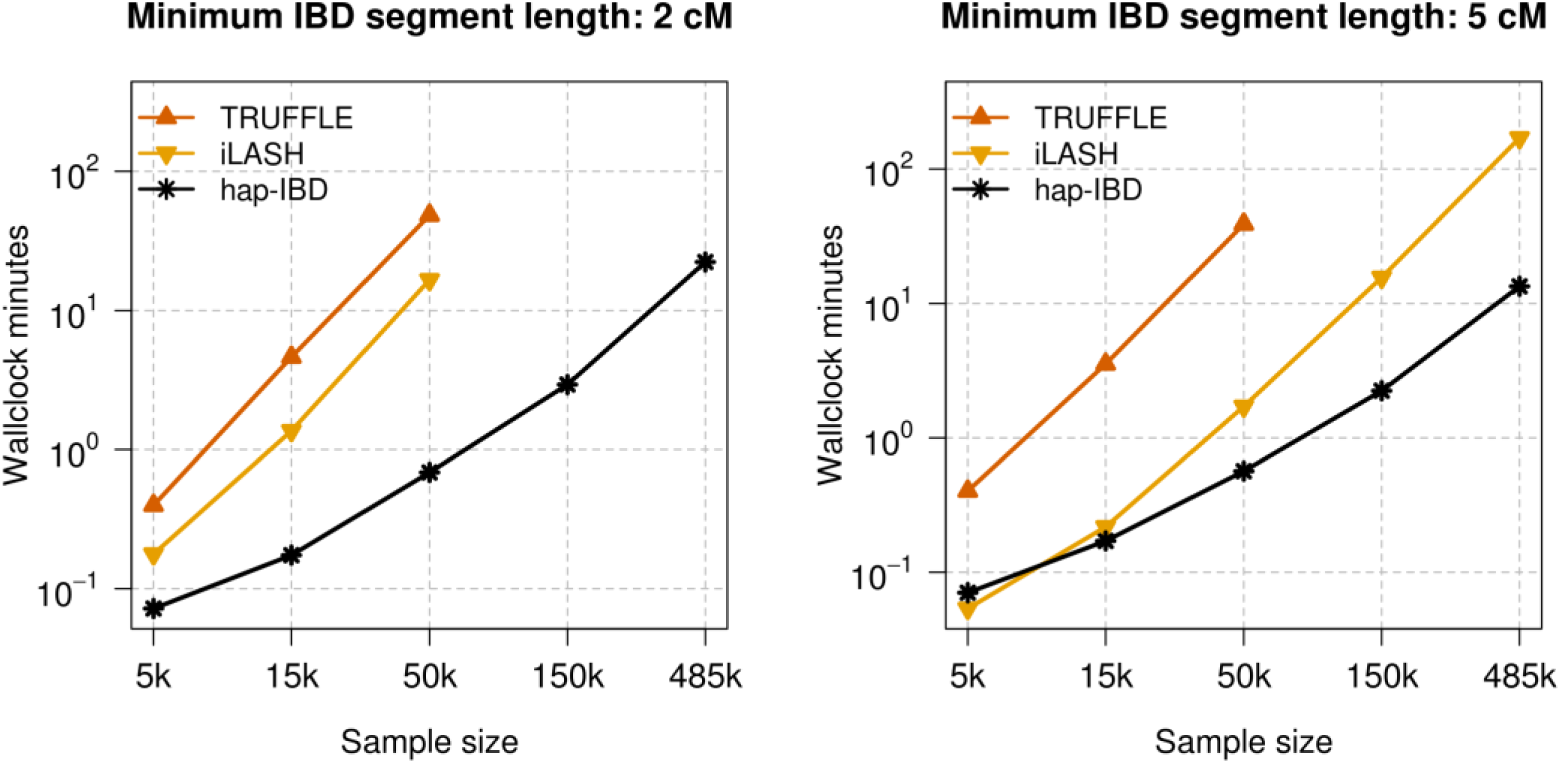
Wall-clock compute time. Wall-clock time for multi-threaded programs when using 12 CPU cores for detecting IBD segments with length ≥ 2 cM (left panel) and ≥ 5 cM (right panel) on chromosome 20 in samples of 5000, 15,000, 50,000, 150,000, and 485,346 individuals from the UK Biobank. All programs used 12 computational threads.

**Figure S2.**
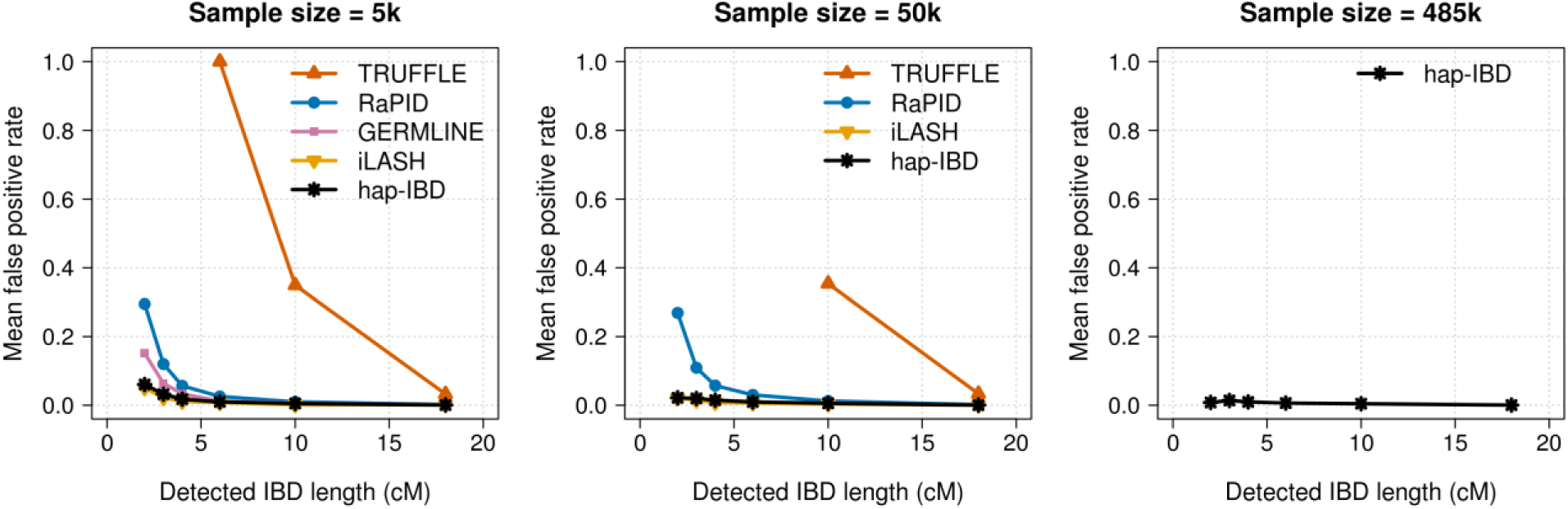
False-positive IBD segment detection in UK Biobank chromosome 20 data. As for Figure 2, but zoomed out to show the full range of false-positive rates. False-positive rates for IBD segment detection for 5000, 50,000, and 485,346 UK Biobank samples. IBD segments with length ≥ 2 cM were detected with each method. Detected IBD segments were assigned into bins of 2-3, 3-4, 4-6, 6-10, 10-18, and >18 cM according to their segment length. The false-positive rate is the proportion of detected IBD segments in a bin that are not covered by any true IBD segment ≥ 1.5 cM in length. Hap-IBD is the only method shown for the full UK Biobank analysis (485,346 individuals) because other methods were unable to complete the analysis with a 2 cM output threshold within the memory and time constraints (see Computational Feasibility Results). The x-coordinate of each data point is the left bin end point (e.g. 2 cM for the 2-3 cM bin).

**Figure S3.**
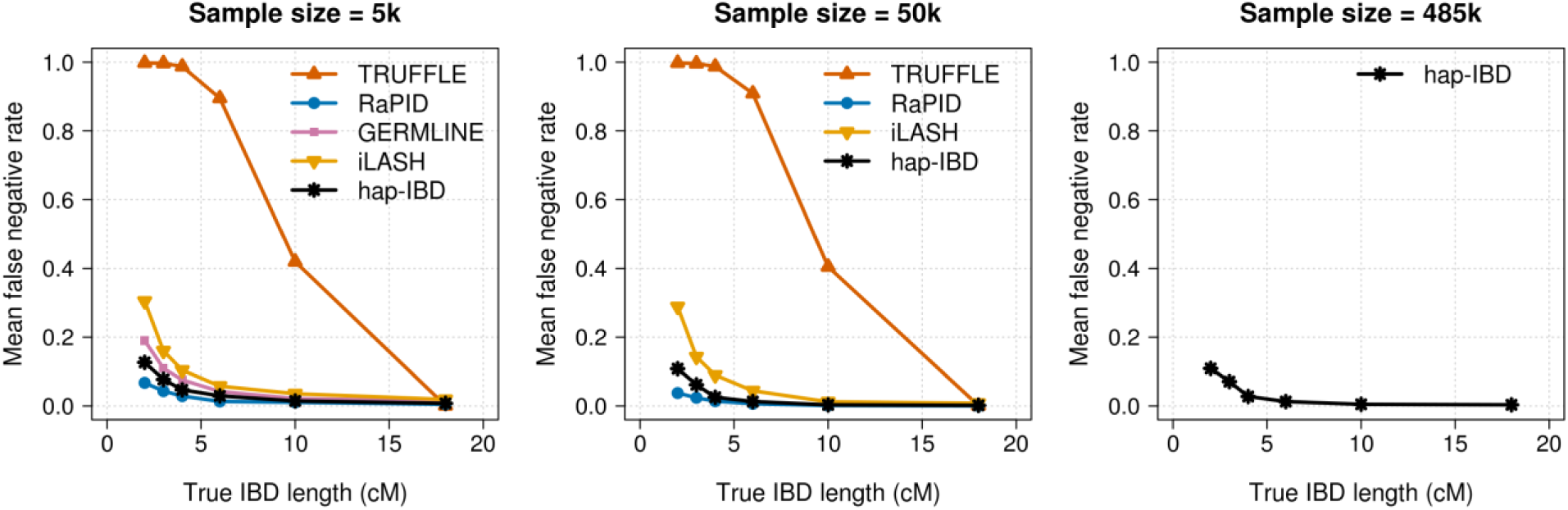
False-negative IBD segment detection in UK Biobank chromosome 20 data. As for Figure 3, but zoomed out to show the full range of false-negative rates. False-negative rates for IBD segment detection for 5000, 50,000, and 485,346 UK Biobank samples. IBD segments with length ≥ 2 cM were detected with each method. The right column of plots shows false negative rates. True IBD segments with length ≥ 2.5 cM were assigned into bins of 2.5-3, 3-4, 4-6, 6-10, 10-18, and >18 cM according to their segment length. The false-negative rate is the proportion of true IBD segments in a bin that are not covered by any detected IBD segment ≥ 2 cM in length. Hap-IBD is the only method shown for the full UK Biobank analysis (485,346 individuals) because other methods were unable to complete the analysis with a 2 cM output threshold within the memory and time constraints (see Computational Feasibility Results). The x-coordinate of each data point is the left bin end point (e.g. 2.5 cM for the 2.5-3 cM bin).

**Figure S4.**
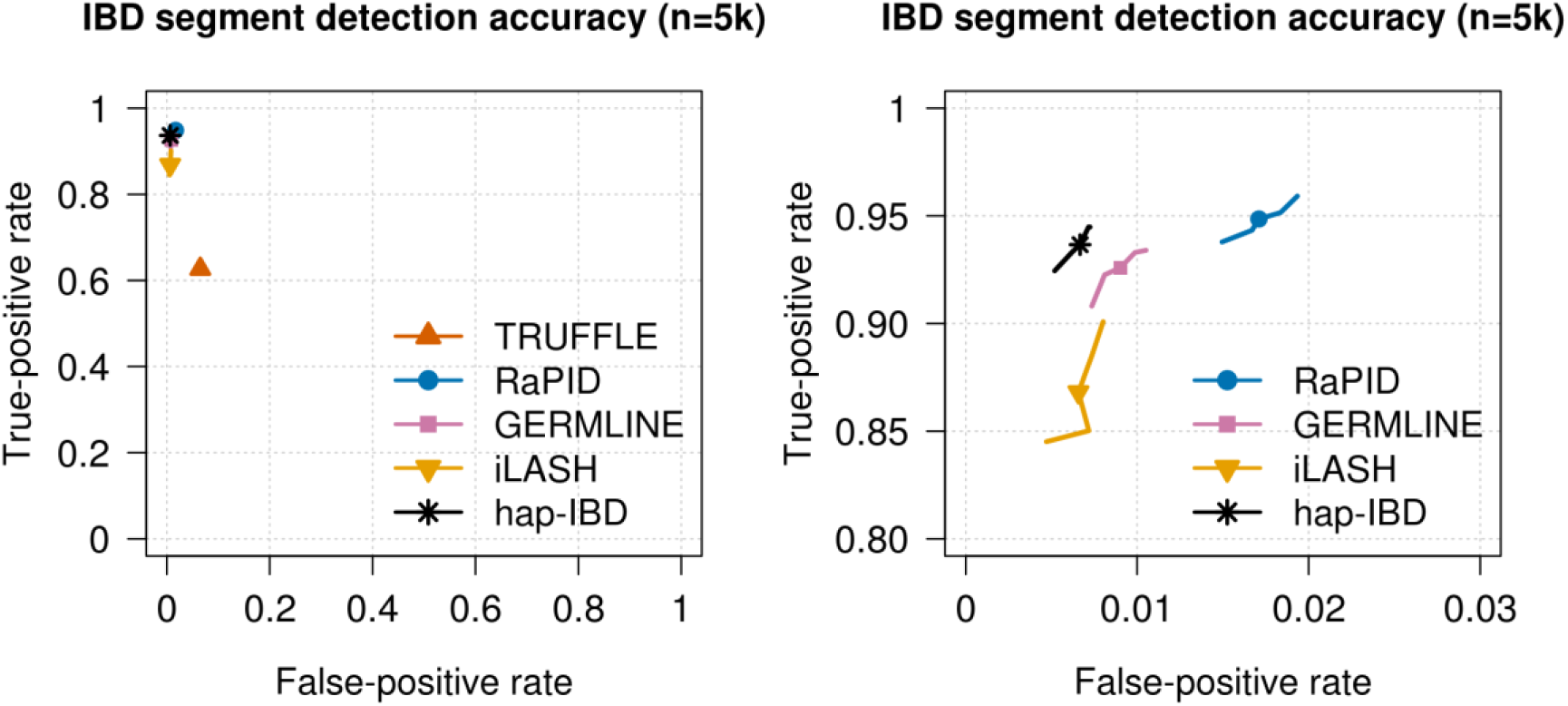
ROC curves for 5 cM IBD segment detection in UK Biobank chromosome 20 data. As for Figure 4, but for 5 cM segments. The right panel is a zoomed in version of the left panel. False-positive and false-negative rates for detection of IBD segments over a range of output length thresholds around 5 cM for 5000 UK Biobank samples. False positives are assessed using true segments of length ≥ 4.5 cM, and false negatives are assessed using true segments of length ≥ 5.5 cM in order to allow for some discrepancy between reported and true lengths. IBD segments were detected with each method using length thresholds of 5 cM (plotted symbol) and with other thresholds between 4.6 and 5.4 cM (lines; see Methods).

**Figure S5:**
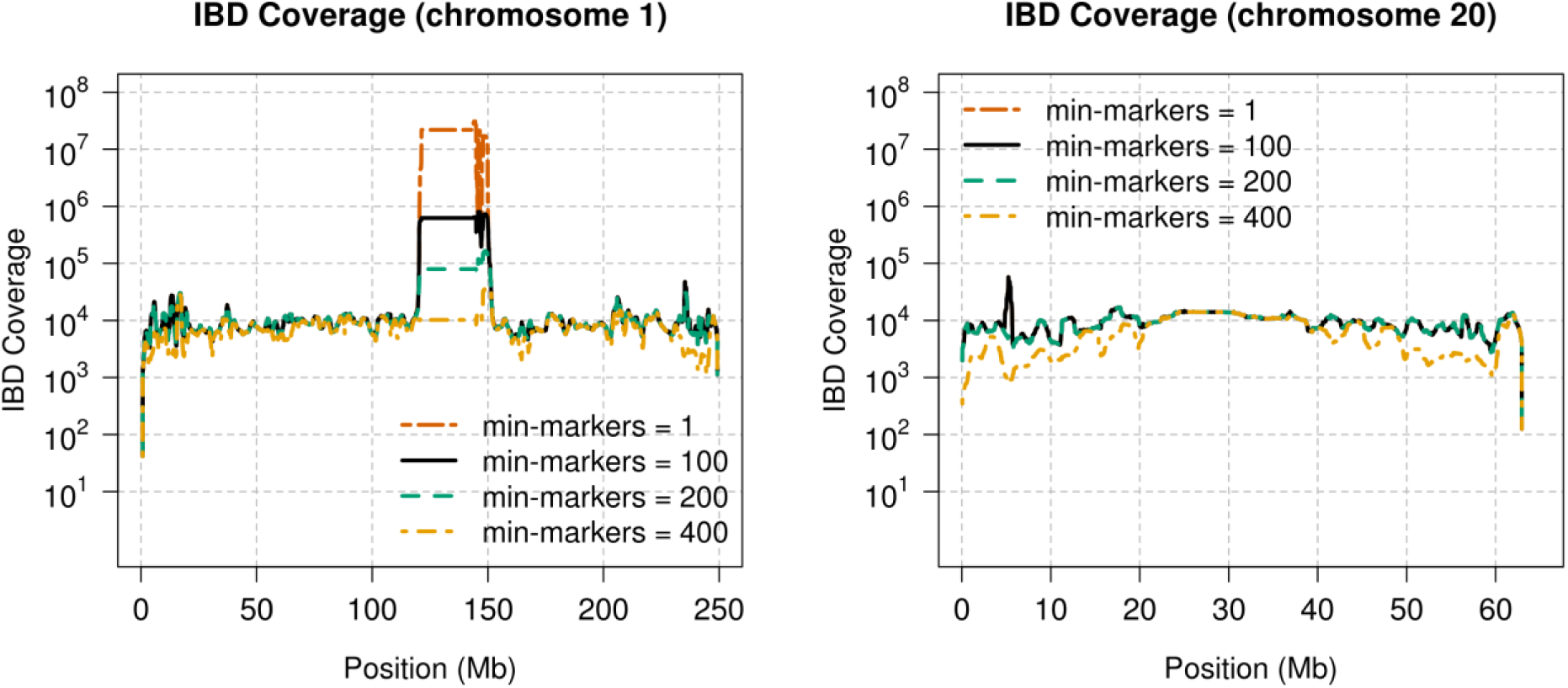
Effect of marker thresholds on IBD segment detection in UK Biobank. The hap-IBD program was run on 5000 UK Biobank samples on chromosome 1 (left panel) and chromosome 20 (right panel) with the min-markers parameter set to 1, 100, 200, and 400, markers. The min-markers parameter controls the minimum number of markers that must be present in a reported seed IBD segment. All other hap-IBD parameters were set at their default values. Each chromosome is divided into non-overlapping 10 kb intervals. For each interval, the IBD segments intersecting the interval are weighted by the proportion of the 10 kb interval that is covered by the IBD segment, and the sum of weights is plotted as the IBD coverage.

**Figure S6.**
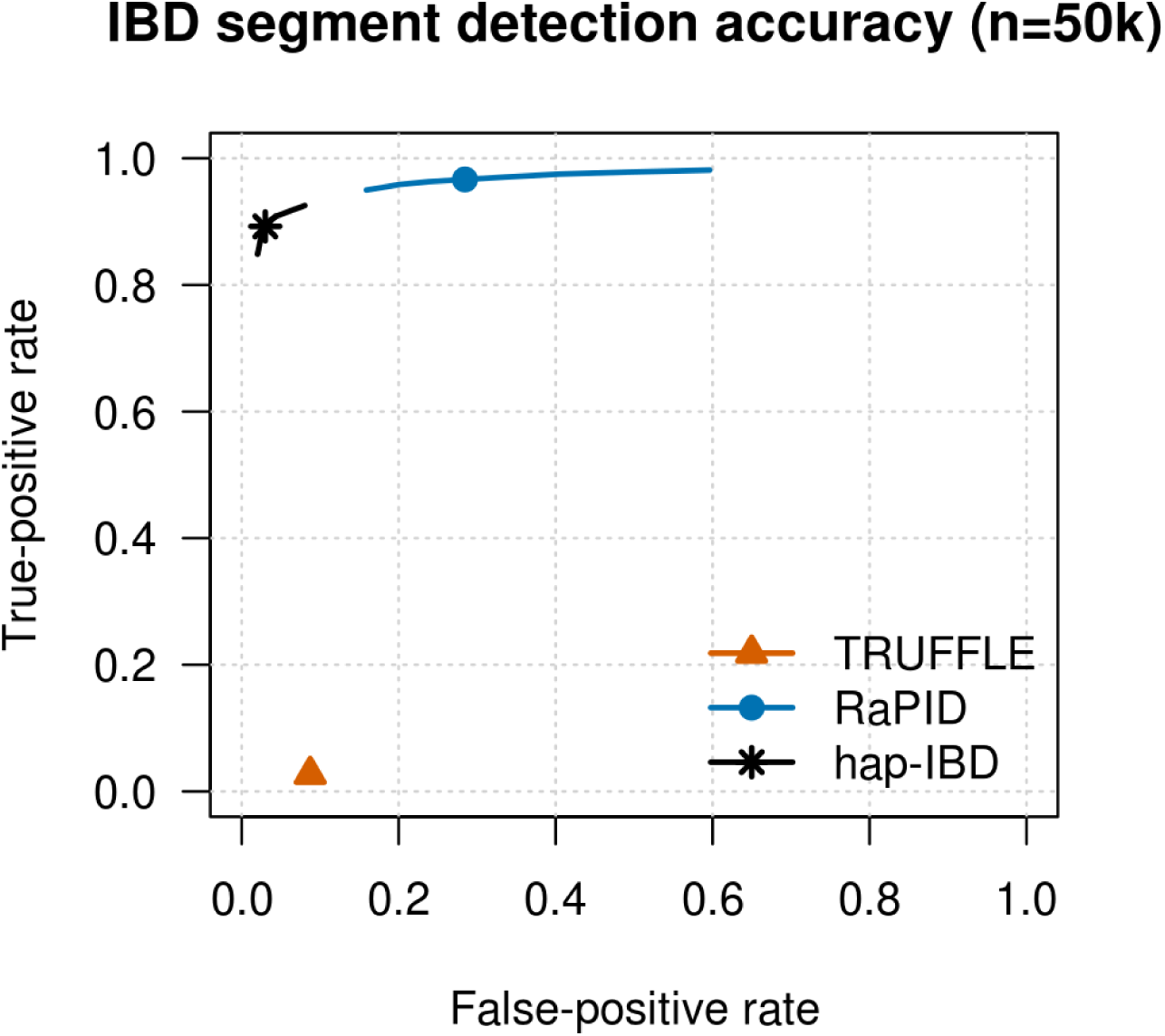
ROC curves for IBD segment detection in simulated sequence data. False-positive and false-negative rates for detection of IBD segments over a range of output length thresholds around 2 cM for 50,000 simulated samples. False positives are assessed using true segments of length ≥ 1.5 cM, and false negatives are assessed using true segments of length ≥ 2.5 cM in order to allow for some discrepancy between reported and true lengths. IBD segments were detected with each method using length thresholds of 2 cM (plotted symbol) and with other thresholds between 1.6 and 2.4 cM (lines; see Methods).

**Table S1:** Parameters used for analysis of UK Biobank data with 2 cM minimum IBD segment length. Parameters that control the output IBD segment length are in red.

| Method | Parameters |
| --- | --- |
| **hap-IBD v1.0** | Default options (min-seed=2.0 min-gap=1000 min-extend=1.0<br>min-output=**2.0** min-mac 2 min-markers=100) |
| **TRUFFLE v1.38** | -mindist 5000 -maf 0.05 -segments -L 1<br>(output was filtered to exclude segments <2.0 cM in length) |
| **GERMLINE 1.5.3** | -haploid -bits 32 -min_m **2.0** |
| **RaPID v1.7** | -r 10 -s 2 -w 3 -d **2.0** |
| **iLASH (commit**<br>**de697321\*)** | perm_count 12 shingle_size 20 shingle_overlap 0<br>bucket_count 4 max_thread 12 match_threshold 0.99<br>interest_threshold 0.7 max_error 0 min_length **2.0**<br>auto_slice 1 cm_overlap 1.4 |

**Table S2:** Parameters used for analysis of simulated sequence data with minimum 2 cM IBD segment length. Parameters that control the output IBD segment length are in red.

| Method | Parameters |
| --- | --- |
| **hap-IBD v1.0** | min-seed=1.0 min-gap=1000 min-extend=0.2 min-output=**2.0** min-markers=100 |
| **TRUFFLE v1.38** | -mindist 5000 -maf 0.1 -segments -L 1<br>(output was filtered to exclude segments <2.0 cM in length) |
| **GERMLINE 1.5.3** | -haploid -bits 75 -min_m **2.0** -err_hom 2 |
| **RaPID v1.7** | -r 10 -s 2 -w 80 -d **2.0** |

